# Mapping Mesoscale Cardiac Architecture with Axially Swept Dithered Light-Sheet Microscopy

**DOI:** 10.1101/2025.10.13.682248

**Authors:** Milad Almasian, Alireza Saberigarakani, Xinyuan Zhang, Elizabeth S. Brewer, Rayaan Hameed, Jichen Chai, Yangyang Lu, Sarah A. Ware, Shuyue Zhou, Yufeng Wen, Hamza Lalami, Hedyeh Hossein Beigi, Jie Yuan, Jun Liao, Kevin M. Dean, Yi Hong, Mingfu Wu, Dan Tong, Darshan Sapkota, Yichen Ding

**Affiliations:** Department of Bioengineering, The University of Texas at Dallas, Richardson, TX, USA; Department of Neuroscience, The University of Texas at Dallas, Richardson, TX, USA; Department of Pharmacological and Pharmaceutical Sciences, University of Houston, Houston, TX, USA; Department of Internal Medicine, Division of Cardiology, UT Southwestern Medical Center, Dallas, TX, USA; Department of Bioengineering, The University of Texas at Arlington, Arlington, TX, USA; Department of Biological Sciences, The University of Texas at Dallas, Richardson, TX, USA; Lyda Hill Department of Bioinformatics, UT Southwestern Medical Center, Dallas, TX, USA; Cecil H. and Ida Green Center for Systems Biology, UT Southwestern Medical Center, Dallas, TX, USA; Center for Imaging and Surgical Innovation, The University of Texas at Dallas, Richardson, TX, USA; Hamon Center for Regenerative Science and Medicine, UT Southwestern Medical Center, Dallas, TX, USA

**Keywords:** microscopy, dithered light-sheet, axially swept approach, volumetric imaging, cardiovascular research, morphological and topological analysis

## Abstract

Comprehensive characterization of cardiac microstructure and vascular networks is critical for assessing cardiovascular development, disease progression, and therapeutic interventions. While volumetric imaging combined with tissue clearing enables high-resolution organ-level analysis, the densely organized cardiac microenvironment, characterized by compacted and trabeculated myocardium, remains technically challenging to resolve in its entirety. To address these limitations, we exploit the temporal dynamics of illumination-detection interplay to create a time-averaged beam profile, thereby maximizing system throughput across multi-scale specimens. Our dithered light-sheet (DiLS) approach extends the confocal region by over 40% and improves the space-bandwidth product while largely preserving optical sectioning capability. Building on this, the axially swept dithered light-sheet (AS-DiLS) employs a dual-actuator strategy that combines rapid beam dithering and axial sweeping to further accelerate imaging speed while achieving micrometer-scale resolution across centimeter-scale scanning ranges, enabling detailed cardiovascular morphological and topological assessment from embryonic to adult stages. We anticipate that this scalable imaging scheme will advance in-depth investigations of organ morphogenesis and disease pathogenesis across a broad spectrum of models.

## 1. Introduction

Cardiovascular diseases remain the leading cause of morbidity and mortality worldwide, with their prevalence projected to rise in the future.[1–3] Cardiac architecture, comprising a complex network of myocardial compaction and trabeculation, vasculature, and extracellular matrix, plays a fundamental role in cardiac function and its adaptability to both physiological and pathophysiological conditions.[4–8] Achieving a comprehensive understanding of organ-level morphology and topology with cellular resolution is critical for elucidating the mechanisms underlying cardiac development, remodeling, protection, and aging, ultimately accelerating the development of effective therapeutic interventions. However, high-fidelity, whole-volume assessment of the microstructure and vascular networks in their entirety within a densely organized cardiac microenvironment, particularly in trabeculated myocardium, still poses substantial technical challenges. In particular, conventional histological and mechanical slicing approaches often compromise tissue integrity and imaging throughput, introducing distortion, tearing, and tissue loss that complicate large-scale reconstructions and generate artifacts that misrepresent intricate microstructures and their spatial relationships within the native in situ environment.[9–13]

While light-sheet microscopy (LSM) has emerged as a viable modality for volumetric imaging with minimal photodamage,[14–17] limitations remain in the mesoscale imaging and analysis of the densely structured three-dimensional (3D) myocardial layers. Specifically, trade-offs in light-sheet thickness, propagation length (e.g., confocal region), and imaging speed remain to be addressed in resolving the volumetric cardiac microstructure and vascular networks across the entire heart.[18–22] Briefly, a thinner Gaussian beam results in better optical sectioning and resolution, but at the cost of a reduced confocal region, limiting the usable field of view (FOV). While the implementation of extended focusing[23] and Bessel beam[24] has improved the effective length of the light-sheet, each presents inherent constraints. Specifically, extended focusing and Bessel beams are accompanied by out-of-focus power that increases with beam length, thereby elevating background and compromising optical sectioning. Other optimized beam configurations, such as Bessel beam with ring patterns on the back focal plane, underutilize the objective’s full aperture and excitation power, thereby limiting their applicability in cardiovascular applications, where dense microstructure, strong scattering, and refractive index heterogeneity exacerbate background accumulation and reduce image contrast across large imaging volumes.[25–27]

To overcome the aforementioned limitations, we conducted a systematic analysis of the temporal coordination between illumination and detection dynamics in current light-sheet imaging systems, with particular emphasis on laser scanning and line intervals inherent to the rolling shutter of sCMOS cameras. This analysis led to the development of an actionable parameter model that identifies key variables and their interdependencies, revealing how confocal region, magnification, exposure time, and line intervals collectively influence optical sectioning, scan range, and image acquisition speed. Building on these insights, we introduce a new method termed dithered light-sheet (DiLS), in which the light-sheet is rapidly oscillated along its propagation direction to generate a time-averaged illumination profile aligned with the subsequently active pixels of the detector. This approach achieves over 40% extension of the confocal region while preserving effective light-sheet thickness compared to a conventional Gaussian beam. Furthermore, DiLS can be seamlessly integrated with axially swept approaches[27–30] by synchronizing the extended focus with the rolling shutter, excluding nonspecific background outside of the effective confocal region. In practical implementation, a voice coil (VC) scanner has been utilized to generate the DiLS, while axial sweeping has been achieved through an electrically tunable lens (ETL) in the prototype, establishing a dual-actuator strategy that integrates rapid beam dithering with axial sweeping to enhance imaging speed while maintaining robust optical sectioning. Our results indicate that the imaging speed of the axially swept dithered light-sheet (AS-DiLS) configuration increases by more than 40% compared to axially swept light-sheet microscopy (ASLM), while maintaining consistent optical sectioning across a scanning range of up to 12.5 mm, with comparable improvements in the space-bandwidth product (FOV / Resolution^2^). This synergy minimizes out-of-focus background while maximizing active pixel utilization and imaging speed. In combination with tissue clearing,[31, 32] AS-DiLS achieves near-isotropic spatial resolution of ∼2.5 µm, enabling mesoscale imaging throughout the trabeculated myocardium, vascular network, and extracellular matrix in intact hearts, effectively addressing photon scattering and optical aberrations caused by heterogeneous refractive indices in muscles.[32–34] Our simulation and experimental results confirm the efficacy of the system design, control configuration, and time-averaged beam profiling across diverse applications.

Collectively, we have demonstrated the capability of the AS-DiLS framework, enabled by a dual-actuator strategy, to capture and analyze the cardiovascular system, including myofibers, extracellular matrix, valves, and vascular trees, addressing longstanding challenges in mesoscale understanding of densely organized myocardium. The morphological and topological assessments enabled by AS-DiLS hold great potential for identifying structural abnormalities associated with both acquired and congenital heart diseases, such as myocardial infarction, hypoplastic left heart syndrome, and outflow tract malformations. In the long term, by linking molecular perturbations to whole-organ architecture, this framework offers a powerful strategy to bridge the gap between genotype and phenotype across multiple organ systems, offering a detailed roadmap and critical structural context to define how specific genetic or signaling disruptions give rise to complex malformations across diverse developmental stages and disease models.

## 2. Results

### 2.1. Standalone DiLS and synchronized AS-DiLS approach

The inherent trade-off between Gaussian beam waist (2*ω*₀) and the extent of the confocal region (**Figure 1A**) imposes limitations on data acquisition speed in axially swept imaging modalities.[27–29] To enhance optical sectioning and accelerate volumetric imaging within the densely structured myocardium of the intact heart, we sought to generate a time-averaged light-sheet strategy by dynamically dithering the beam waist along its propagation direction. The effective light-sheet thickness is typically accepted as up to 2√2*ω*₀ for a Gaussian beam profile.[14, 35] In this case, we used a VC scanner integrated with a remote focusing setup[36, 37] to dither the beam waist along its propagation axis at a frequency set to twice the reciprocal of the camera exposure time (e.g., dithering frequency *f_VC_* = 2/*ET*, where *ET* is exposure time) (**Figures 1B**, **S1**). Beam profiles of DiLS with varying dithering ranges were captured using 50 μM fluorescein under a 20 ms exposure time and a dithering frequency of 100 Hz (**Figure 1C**).

**Figure 1.**
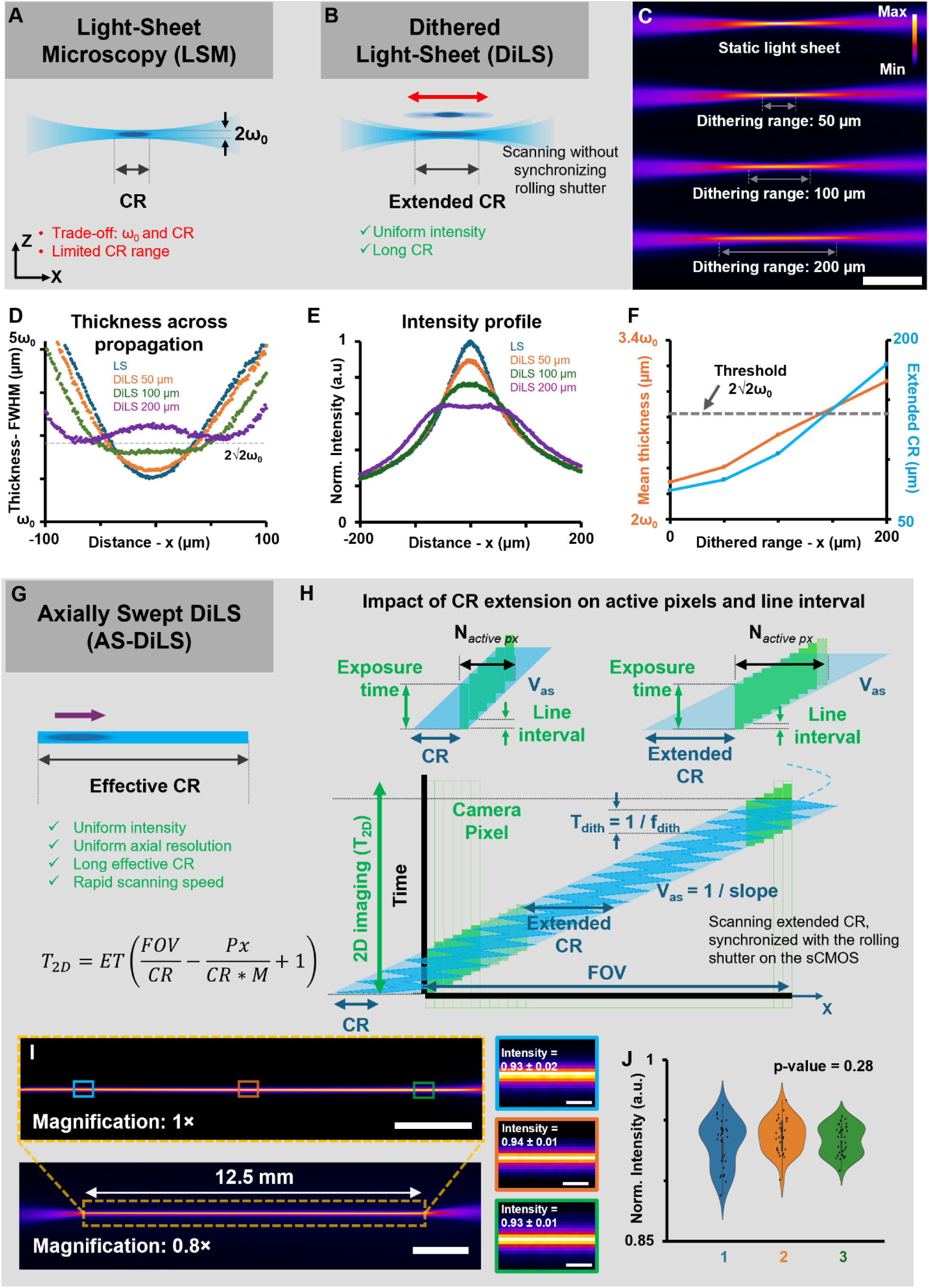
Dithered and axially swept dithered light-sheet microscopy. **(A)** Conventional LSM schematic showing the trade-off between light-sheet thickness (2*ω*₀) and confocal region (CR). **(B)** DiLS extends the CR by dithering the light-sheet along its propagation direction, generating a time-averaged beam. **(C)** Experimental profiles of static and dithered light-sheets with 50, 100, and 200 µm ranges in 50 µM fluorescein solution. **(D)** Thickness variations along the propagation axis quantified by FWHM. **(E)** Normalized intensity profiles as a function of dithering range. **(F)** Quantification of average light-sheet thickness in the CR (orange) and the extended CR (blue) as functions of the dithering range, showing optimal extension below the 2√ 2*ω*₀ threshold. **(G)** Concept of AS-DiLS integrating DiLS with axial sweeping to cover the full imaging plane. **(H)** Scanning mechanism with the extended CR synchronized to the rolling shutter of the sCMOS camera. **(I)** Illumination profile in fluorescein solution, sampled at left, middle, and right regions, each spanning 195 µm (data are presented as mean ± SD). **(J)** Quantification of normalized intensity across three positions (one-way ANOVA, *p* = 0.28, n = 30 measurements per region), confirming uniform distribution across the imaging plane. Px, camera pixel size; M, magnification; FOV, field of view; ET, exposure time. Scale bars: **(C)** 100 µm, **(I)** 2 mm and 50 µm (insets).

The results presented at dithering ranges of 0, 50, 100, and 200 µm revealed notable changes in effective light-sheet thickness (**Figure 1D**), spatial intensity distribution (**Figure 1E**), and axial extent of the confocal region (**Figure 1F**). Using our prototype setup (Plan Fluor 10×/0.3 NA, Nikon), we established a baseline confocal region of 73 µm. To balance thin light-sheet with extended axial coverage, a 100 µm dithering range was identified as optimal in practice (**Figure 1F**). Under this condition, the effective light-sheet thickness remained below the threshold of 2√2*ω*₀, varying from 2.57*ω*₀ to 2√2*ω*₀ (10.1% relative range), while the corresponding confocal region extended to approximately 105 µm, a 44% improvement over the baseline (**Figure 1D**). In contrast, a 50 µm dithering range yielded a narrower confocal region of 83 µm and a higher thickness relative range (29.7%, 2.18*ω*₀ to 2√2*ω*₀), while a 200 µm range resulted in a beam thicker than 2 √2*ω*₀, falling outside the preferred region, despite improved intensity uniformity as the dithering range increases (**Figure 1E**). This condition represents a practical balance between extending the confocal region and preserving optical sectioning, rather than eliminating the inherent trade-off between light-sheet thickness and propagation length. Simulation results further supported the experimental findings that a 100 µm dithering range is basically optimal (**Figure S2** and **Note S1**), and this configuration was adopted for subsequent cardiovascular applications.

To extend the imaging coverage up to centimeter scales, DiLS is able to be integrated with axially swept approaches,[27–29] enabling a new mode termed AS-DiLS (**Figure S1**). In practice, an ETL was added as a secondary scanner into the prototype to axially sweep the dithered beam across the imaging plane, ensuring tight synchronization between the effective confocal region of DiLS and the active pixels of the sCMOS camera (**Figure 1G**). In AS-DiLS, the camera’s exposure time is used to determine the dithering frequency, and the 2D image acquisition time (*T_2D_*) is used to derive the axial sweeping frequency (*f_ETL_* = 1/(2×*T_2D_*)) (**Note S2** and **Equations S1-4**). Briefly, selective pixels (green grid in **Figure 1H**) are activated upon arrival of the extended beam waist and remain active throughout the exposure period (solid green area), during which they collect signal exclusively from the confocal region (dark blue area). In DiLS, the beam waist dithers back and forth with a period equal to half the exposure time, thereby generating an extended confocal region (light blue area). Compared to LSM with the same exposure time, this extended confocal region in DiLS led to an increased number of active pixels and consequently reduced the line interval time between successive pixel activations (**Note S2** and **Equations S9, 10**), which is an inherent limitation in conventional axially swept modalities. By synchronizing the rolling shutter with the extended beam waist, the reduced acquisition time to generate a 2D frame is defined as follows:

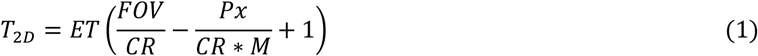

where *T_2D_* is the acquisition time of an image, *ET* is the exposure time, *FOV* denotes the field of view, *CR* represents the confocal region, *Px* refers to the pixel size of the camera sensor, and *M* is the magnification (**Note S3** and **Equations S11, 12**). Using a 100 µm dithering range, we demonstrated over 40% improvement in imaging speed in AS-DiLS (**Figure 1H** and **Note S3**). Our fluorescein experiments indicated that AS-DiLS achieved a maximum scanning range of 12.5 mm under 1× magnification (**Figure 1I**). Spatial intensity analysis at the left (0.93 ± 0.02 a.u.), middle (0.94 ± 0.01 a.u.), and right (0.93 ± 0.01 a.u.) regions of the beam showed no statistically significant differences (*p* = 0.28), indicating uniform beam distribution across the scanning field (**Figure 1J**).

### 2.2. Characterization of the system

We experimentally characterized system performance under four configurations, LSM, ASLM, DiLS, and AS-DiLS, all implemented into a single prototype system equipped with a VC scanner and an ETL[38] (**Figure S1**). A customized LabVIEW program was developed to synchronize device operations, including the sCMOS camera, VC scanner, and ETL. Additionally, an executable MATLAB script, customized to automate the setup of calculation parameters, including axially swept frequency (*f_ETL_*), dithering frequency (*f_VC_*), ETL amplitude, and line interval, has been shared in Zenodo.[39] It ensures flexible operation and reproducibility across various experimental conditions (**Figure S4**).

The light-sheet beam generated by the cylindrical lens was dithered using the VC scanner at frequencies ranging from 20 to 100 Hz in DiLS, while the extended focus was axially swept using the ETL at frequencies between 0.25 and 25 Hz in AS-DiLS. The detection module, consisting of a 1×/0.25 NA objective lens (MV PLAPO, Olympus) and a zoom body (0.63–6.3×, MVX-ZB10, Olympus), provides a long working distance of 87 mm and a flexible FOV from 21 mm to 2.1 mm. This design addresses constraints associated with working distance for intact rodent hearts, reduces challenges posed by refractive index matching solutions, streamlines multi-immersion protocols for tissue-cleared samples within the imaging chamber, and leverages the extended depth of field via spherical aberration correction.[40]

Spatial resolution was assessed using 0.53 µm fluorescent beads. Under the LSM configuration, the minimum axial resolution at the beam waist was 2.24 µm. Consequently, the confocal region, defined as the range where axial resolution remains below 3.15 µm was measured to be 73 µm. The lateral resolutions (in blue) of three representative beads located to the left, within, and to the right of the confocal region were 2.31 µm, 2.25 µm, and 2.11 µm, respectively, while the axial resolutions (in red) were 6.09 µm, 2.69 µm, and 6.87 µm, respectively, indicating the varying light-sheet thickness along the propagation direction (**Figure 2A** and data for all beads in **Figures S5A, B**). Within the confocal region, the lateral resolution of all beads (n = 103) was 2.25 ± 0.08 µm along the x-axis and 2.79 ± 0.13 µm along the y-axis, with an average axial resolution of 2.63 ± 0.30 µm.

**Figure 2.**
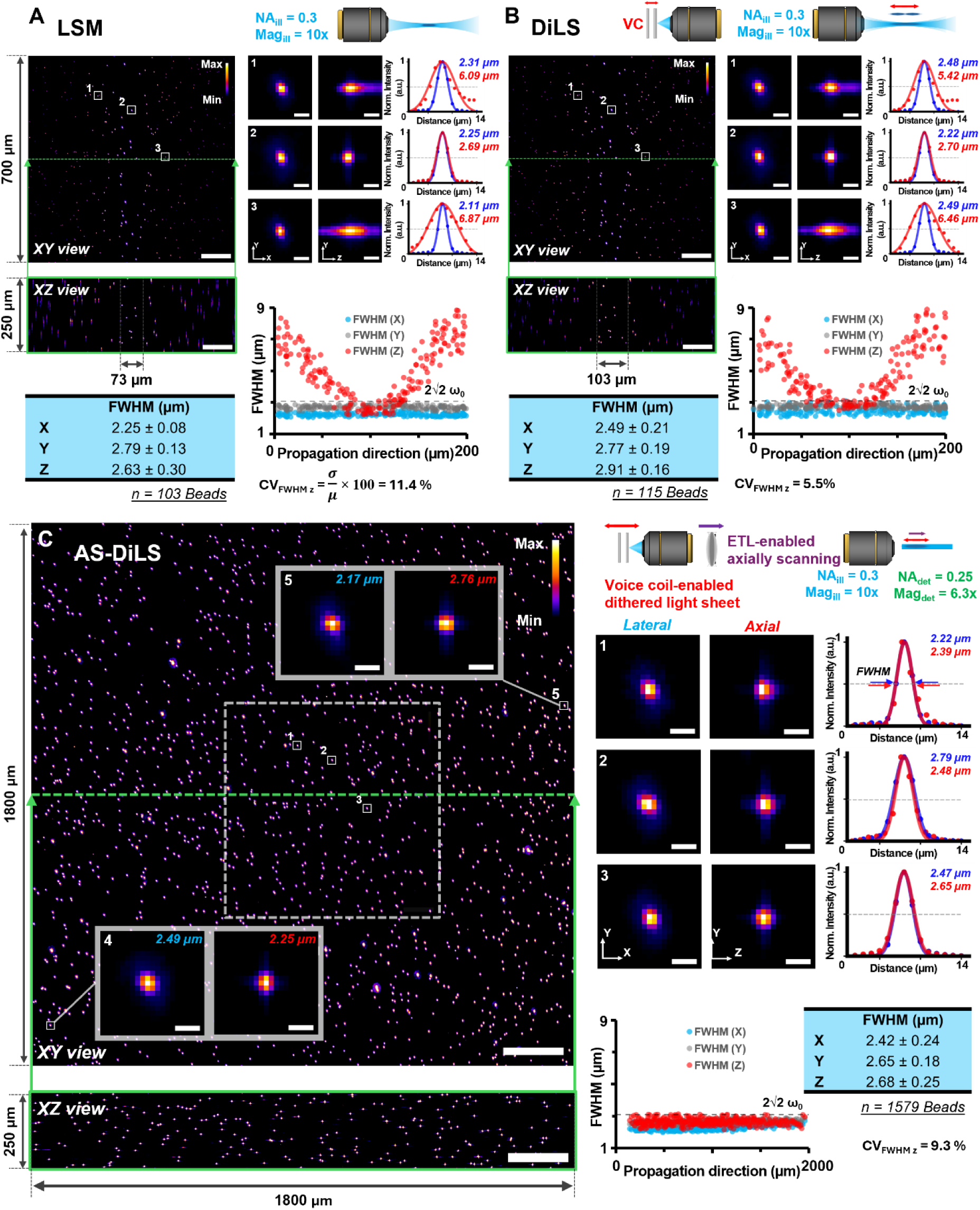
Comparative analysis of imaging performance in LSM, DiLS, and AS-DiLS. **Volumetric** imaging of 0.53 µm fluorescent beads in 1% agarose gel using conventional LSM **(A)**, DiLS **(B),** and AS-DiLS **(C)**. Cropped volumes of 700 × 700 × 250 µm^3^ shown for LSM and DiLS; dashed box in **(C)** marks the crop region. Maximum intensity projections (MIP) of XY and XZ views displayed in Fire pseudo color. Insets show selected beads at different positions used for resolution analysis. Three beads located at different positions were selected for lateral and axial resolution analysis for LSM, DiLS, and AS-DiLS. Lateral (blue) and axial (red) intensity profiles were fit with Gaussian distributions, and FWHM values calculated. For LSM and DiLS, beads within the usable confocal region were analyzed; for AS-DiLS, resolved beads across the swept imaging field were analyzed (n = 103 for LSM, n = 115 for DiLS, and n = 1579 for AS-DiLS). The average FWHM and standard deviation along the X, Y, and Z axes were computed and tabulated. In AS-DiLS, beads at the far left and right of the 1800 × 1800 µm^2^ image were also analyzed. For bead-based point spread function (PSF) analysis, n indicates the number of resolved beads analyzed as technical measurements. Scale bars: 200 µm (XY and XZ views) and 5 µm (bead insets).

Under DiLS conditions with *ET* = 20 ms and *f_VC_* = 100 Hz, the lateral resolutions of the same representative beads were 2.48 µm, 2.22 µm, and 2.49 µm, respectively, while the axial resolutions were 5.42 µm, 2.22 µm, and 6.46 µm, respectively (**Figure 2B** and data for all beads in **Figures S5C, D**). Using the same threshold of 3.15 µm as in LSM, the extended confocal region was determined to be 103 µm (41% improvement). Within this region, the mean lateral resolutions (n = 115) were 2.49 ± 0.32 µm along the x-axis and 2.77 ± 0.19 µm along the y-axis, with an average axial resolution of 2.91 ± 0.16 µm. Thus, standalone DiLS extended the usable confocal region from 73 µm to 103 µm, while resulting in a modest increase in the mean axial FWHM from 2.63 µm to 2.91 µm. This result indicates a practical balance between confocal-region extension and optical sectioning, rather than complete elimination of the inherent trade-off between light-sheet thickness and propagation length (**Figure S6**). In comparison, AS-DiLS yielded lateral resolutions of 2.22 µm, 2.79 µm, and 2.47 µm, and axial resolutions of 2.39 µm, 2.48 µm, and 2.65 µm. Measurements of two additional beads located at the extreme left and right of the FOV indicated lateral resolutions of 2.49 µm and 2.17 µm, and axial resolutions of 2.25 µm and 2.76 µm, respectively. Comprehensive analysis of all resolved beads (n = 1579) captured by AS-DiLS revealed lateral resolutions of 2.42 ± 0.24 µm (x-axis) and 2.65 ± 0.18 µm (y-axis), and an axial resolution of 2.68 ± 0.25 µm, across 1.8 mm at 6.3× zoom magnification (**Figure 2C**). Importantly, AS-DiLS maintained this near-isotropic resolution across an FOV that was 25 times larger than that of conventional LSM. As a result, the space-bandwidth product, a qualitative metric for assessing an image’s information-carrying capacity, is therefore enhanced.[41] These results demonstrate that AS-DiLS achieves near-isotropic resolution and consistent optical sectioning throughout the entire scanned range. In addition, the coefficient of variation (CV) was utilized to quantify the relative variability in axial resolution within each imaging configuration. Our results showed that CV values of 11% for LSM (n = 103), 5.5% for DiLS (n = 115), and 9.3% for AS-DiLS (n = 1579), with bootstrapped 95% confidence intervals of 10.3%–12.1%, 4.8%–6.0%, and 9.3%–9.8%, respectively.

These results indicate low relative axial-resolution variability within the usable confocal regions of LSM and DiLS and across the larger swept imaging field of AS-DiLS. The LSM and DiLS CV values were calculated from beads located within their respective usable confocal regions, whereas the AS-DiLS CV value was calculated from beads distributed across the larger swept imaging field. Therefore, the CV values were interpreted as configuration-specific descriptors of axial-resolution uniformity.

### 2.3. Comparison of AS-DiLS with ASLM

To evaluate the effect of replacing the static light-sheet used in ASLM with a DiLS under an otherwise identical axially swept, rolling-shutter synchronized acquisition, we compared ASLM (generated by static light-sheet) and AS-DiLS (generated by DiLS) across imaging speeds from 1 to 25 Hz (**Figure 3A**). In the non-sweeping configuration, the measured sheet thickness (2*ω*₀) was 3.13 µm for static light-sheet and 3.61 µm for DiLS (**Figure 3A**). After enabling axial sweeping with rolling-shutter synchronization, the effective light-sheet thickness increased with imaging speed for both methods; however, AS-DiLS sustained a thinner sheet at higher speeds, owing to its synchronization architecture, which tolerates larger timing deviations among key components. Specifically, ASLM increased from 3.25 ± 0.07 µm at 1 Hz to 8.27 ± 1.44 µm at 25 Hz (154.5% increase), whereas AS-DiLS increased from 3.63 ± 0.09 µm to 7.83 ± 1.34 µm (115.7% increase) over the same range, yielding significantly thinner light-sheets for AS-DiLS at speeds above 2 Hz (Wilcoxon signed-rank test, *p* < 0.001; **Figure 3B**). At low imaging speeds (≤ 2 Hz), ASLM produced slightly higher median normalized intensity than AS-DiLS (0.968 vs 0.934 a.u. at 1 Hz; 0.829 vs 0.814 a.u. at 2 Hz). In contrast, at acquisition rates above 2 Hz, AS-DiLS yielded higher intensity, with an increase of up to 19.2% at 5 Hz (0.562 vs 0.454 a.u.), and produced significantly higher intensity than ASLM at 10, 15, 20, and 25 Hz (Wilcoxon signed-rank test, *p* < 0.001; **Figure 3C**).

**Figure 3.**
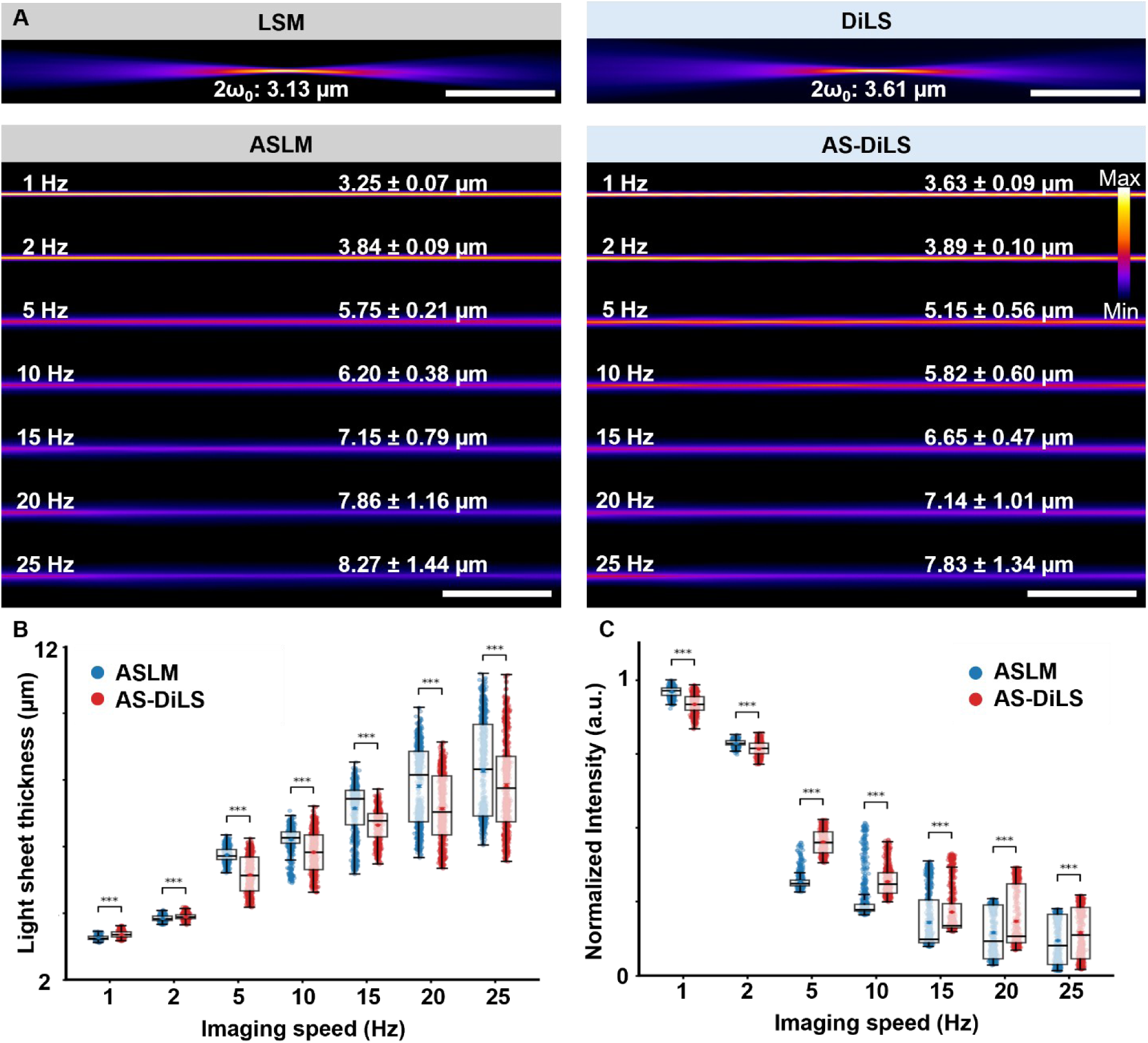
Comparison of AS-DiLS and ASLM. **(A)** Representative fluorescein intensity profiles for a static light-sheet (LS) and a dithered light-sheet (DiLS), each implemented in the same axially swept, rolling-shutter synchronized configuration to form ASLM and AS-DiLS, over imaging speeds of 1–25 Hz. Data are presented as mean ± SD of light sheet thickness. **(B)** Comparison of effective light-sheet thickness as a function of imaging speed for ASLM and AS-DiLS. Thickness increased with speed for both methods; AS-DiLS remained significantly thinner at speeds >2 Hz (n = 851 measurements for each group; Wilcoxon signed-rank test, *** *p* < 0.001). **(C)** Comparison of mean normalized intensity versus imaging speed for ASLM and AS-DiLS (n = 851 measurements for each group; Wilcoxon signed-rank test, *** *p* < 0.001). *P* values in B and C were calculated using a Wilcoxon signed-rank test. Scale bars: **(A)** 200 µm.

### 2.4. Assessment of image quality in hearts

Significant light attenuation primarily caused by fibrous tissue and heterogeneous refractive indices within the densely structured myocardium consistently degrades the image quality in conventional volumetric imaging of intact hearts. Our methodology is compatible with multiple established tissue clearing methods, most notably iDISCO for demonstration in this study.[31, 42] Prior to imaging, cardiac specimens were cleared and embedded in 1% agarose gel and filled with the refractive index-matching solution for imaging by either autofluorescence or immunolabeling (**Figures 4A, B**). It should be noted that iDISCO-based clearing may lead to tissue deformation.[32, 43] To evaluate the consistency of structural information across the entire imaging plane in AS-DiLS, a cleared mouse heart was translated laterally in six increments of 341 µm, with an image acquired at each position under the same imaging conditions (**Figure 4C**). The selected region of interest (341 × 2048 µm) was assessed across six sequential images using the structural similarity index measure (SSIM). With values ranging from 0.96 to 0.99, the analysis revealed a high degree of similarity between adjacent regions, indicating that the optical sectioning capability of AS-DiLS is spatially independent and effectively preserves physiological structural information (**Figure 4D**).

**Figure 4.**
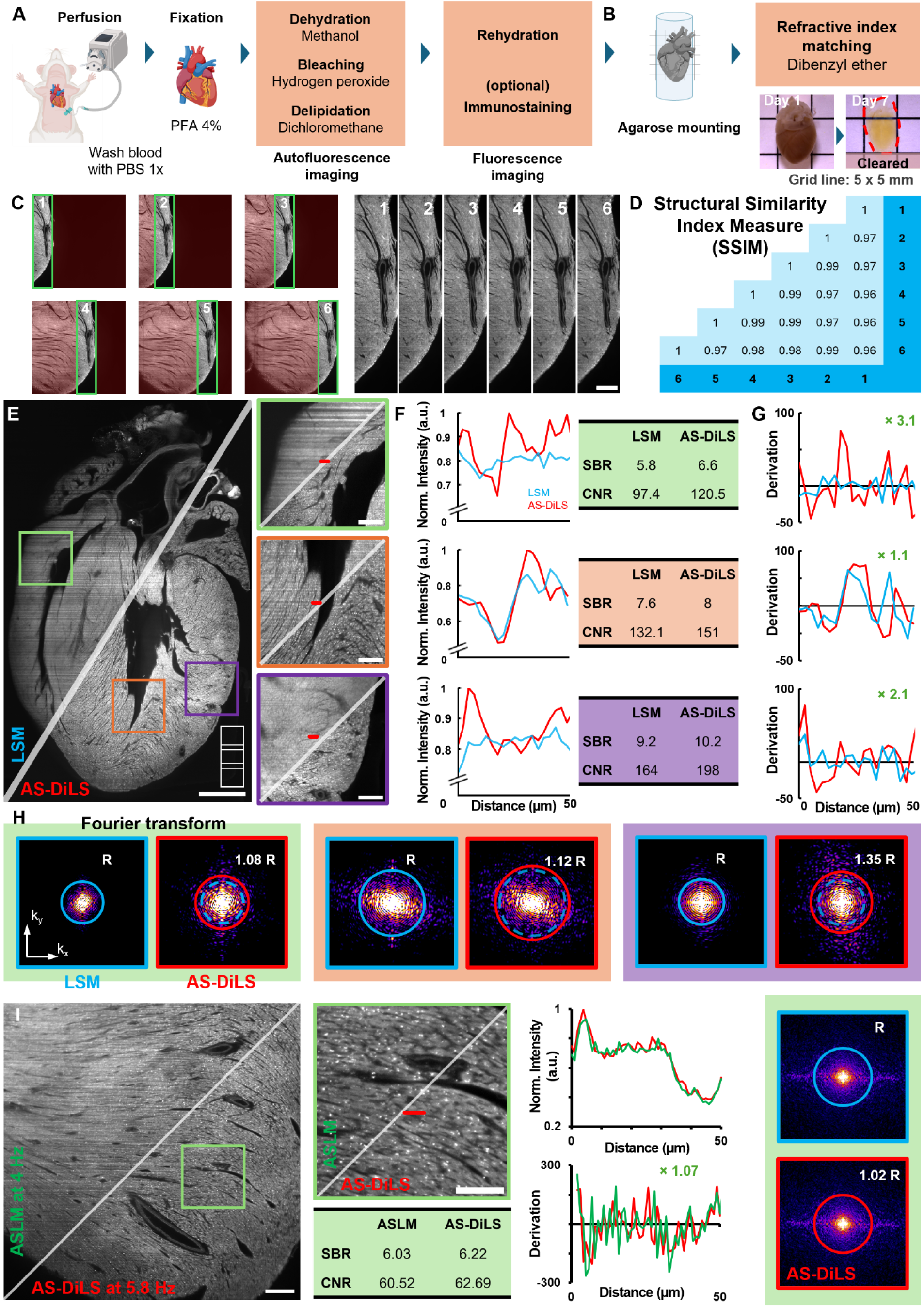
Mouse heart clearing and quantitative evaluation of AS-DiLS images. **(A)** Schematic of iDISCO-based clearing protocol adapted for autofluorescence and immunofluorescence imaging of intact mouse hearts. **(B)** Heart embedding in agarose and subsequent clearing in refractive index matching. **(C)** Imaging uniformity assessed by acquiring the same region of the heart in six adjacent lateral sections across the FOV. **(D)** SSIM values between adjacent tiles ranged from 0.96 to 0.99, confirming consistent performance across the imaging plane. **(E)** Comparison of image quality in 2D cross section between LSM (left) and AS-DiLS (right). Three representative 1 × 1 mm regions (green, orange, and purple boxes) from the left, middle, and right portions of the heart were selected for quantitative analysis. **(F)** Normalized intensity profiles for each region, with SBR and CNR values reported. **(G)** First-derivative plots of intensity profiles comparing LSM and AS-DiLS showed sharper edges with AS-DiLS. **(H)** Fourier transform analysis of representative regions revealed a larger cut-off radius in AS-DiLS, reflecting improved support of lower-bound spatial frequencies. **(I)** Imaging speed comparison under matched image quality conditions in representative cross-sections of the adult mouse heart, acquired using ASLM at 4 Hz and AS-DiLS at 5.8 Hz. Quantitative metrics include SBR, CNR, edge sharpness derived from the first derivative of line intensity profiles, and Fourier analysis of spatial frequency support based on OTF cut-off radius. All quantitative analyses were performed using raw 16-bit image data. Grid line in **(B)** 5 mm. Scale bars: **(C)** 200 µm, **(E)** 1 mm and 200 µm (insets), and **(I)** 500 µm and 100 µm (insets).

To further assess AS-DiLS performance in the heterogeneous cardiac tissues, three representative regions (1 × 1 mm) of the mouse heart were selected along the light-sheet propagation axis: the distal (green), the confocal region (orange), and the proximal ends (purple) (**Figure 4E** and **Table S1**). AS-DiLS consistently revealed the myocardial microstructures across three sections spanning over 5 mm, in contrast to the degraded image quality at both ends when AS-DiLS was disabled (**Video S1**). All quantitative analyses were performed using raw 16-bit image data. Quantitative analysis indicated that activating AS-DiLS increased the signal-to-background ratio (SBR) by 13.8% (from 5.8 to 6.6), 5.3% (from 7.6 to 8.0), and 10.9% (from 9.2 to 10.2) in the distal, confocal, and proximal regions, respectively. Similarly, contrast-to-noise ratio (CNR) improved by 23.7% (from 97.4 to 120.5), 14.3% (from 132.1 to 151.0), and 20.7% (from 164.0 to 198.0) across the same regions, affirming robust signal and contrast enhancement throughout heterogeneous, densely structured myocardial tissue using AS-DiLS (**Figure 4F**). Image sharpness, assessed by the first derivative of intensity profiles, increased 2.1-fold and 3.1-fold at the distal and proximal ends, respectively, while remaining ∼1.1-fold within the confocal region (**Figure 4G**), demonstrating improved optical sectioning capability outside the confocal region with AS-DiLS. Our Fourier analysis further revealed the optical transfer function (OTF) cut-off radius, which represents the lower-bound estimate of supported frequencies,[44] increased by 8%, 12%, and 35% in the distal, confocal, and proximal regions, respectively, indicating improved capture of fine structural details enabled by AS-DiLS (**Figures 4H, S11**). To compare AS-DiLS and ASLM at matched image quality, representative cross-sections of the adult mouse heart were imaged using AS-DiLS at 5.8 Hz and ASLM at 4 Hz. Under comparable SBR (6.22 vs 6.03), CNR (62.69 vs 60.52), edge sharpness, and OTF cut-off radius, AS-DiLS enabled a 45% faster acquisition rate without compromising image quality relative to ASLM (**Figure 4I**).

### 2.5. Morphological analysis of the cardiac specimens

To demonstrate whole-heart imaging capability, we implemented AS-DiLS into the imaging of myocardium from mouse embryos to neonates to adults. To cover an intact adult mouse (14 months, male, C57BL, The Jackson Lab), we stitched three consecutive AS-DiLS tiles from the original 16-bit raw image data, with a 10% overlap along the ventricular long axis. The entire image stack includes a volumetric region measuring 5.3 × 9 × 5 mm^3^, with an acquisition time of approximately 30 min per tile for ∼1000 images, and a total data size of 13.3 GB that remarkably improves information density for fine morphological details in AS-DiLS (**Figure 5A** and **Table S1**).

**Figure 5.**
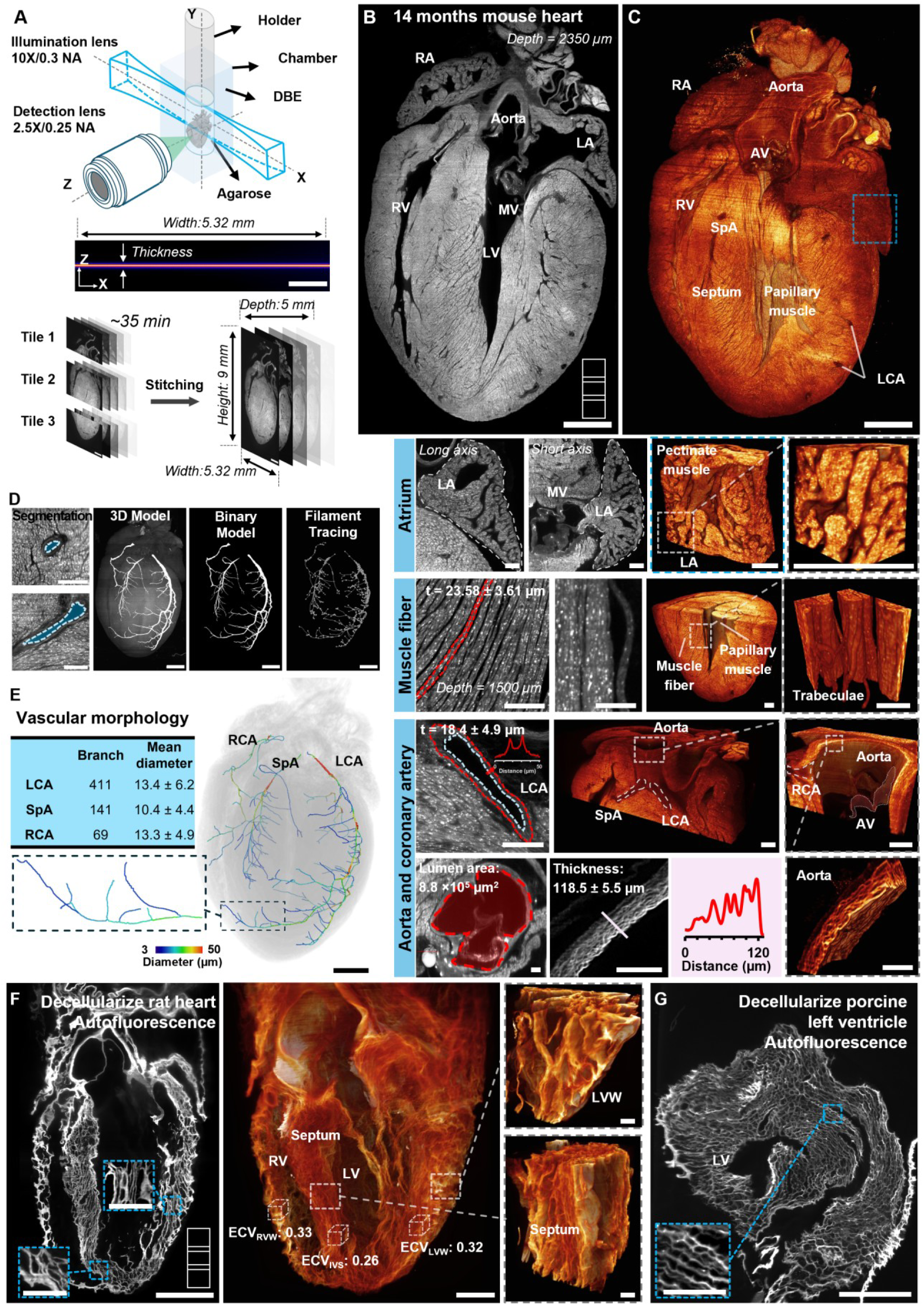
AS-DiLS autofluorescence imaging and in-depth morphological analysis. **(A)** Schematic representation of the light-sheet imaging setup for intact adult heart volumetric imaging. **(B)** Representative 2D cross-section at a depth of 2350 µm. **(C)** 3D volumetric reconstruction processed with zoomed views resolving trabeculae, fibers, and vessel morphology. **(D)** Pipeline for tracing and segmentation of coronary arteries. **(E)** Quantitative vascular morphology of the coronary artery network, including branch count and lumen diameter, with a color-coded diameter map. For vessel diameter analysis, n represents the number of vessel branches analyzed for each coronary artery: LCA, n = 411; SpA, n = 141; RCA, n = 69. Data are presented as mean ± SD. **(F)** Volumetric reconstruction of a decellularized adult rat heart imaged under 2× magnification using a tiled acquisition. Autofluorescence from extracellular matrix enables label-free visualization of structures throughout the tissue. **(G)** AS-DiLS imaging of extracellular matrix in a decellularized left ventricle of a porcine heart. Vessel diameter and branch measurements were obtained from multiple vessel segments within the same representative adult heart. RA, right atrium; LA, left atrium; RV, right ventricle; LV, left ventricle; MV, mitral valve; AV, aortic valve; RCA, right coronary artery; LCA, left coronary artery; SpA, septal artery. Scale bars: 1 mm and 200 µm (insets).

The representative myocardial section along the long axis (depth = 2350 µm) (**Figure 5B** and **Video S2**) and the 3D rendering (**Figure 5C**) provided an entry point to investigate myocardial architecture at both the global pattern and regional detail levels. These include the pectinate muscle in atria, myocardial fibers, papillary muscle, and trabecular network in ventricles, aortic valve leaflet associated with the ascending aorta, and coronary arteries throughout the heart (Insets in **Figure 5C** and **Videos S3,4**). The near-isotropic resolution enables quantitative analysis of well-resolved microstructure from physiologically relevant perspectives, such as long- and short-axis views, after volumetric rendering. For instance, our results revealed that the average thickness of a representative myocardial fiber, measured at 10 points, was 23.6 ± 3.6 µm (**Figure 5C, middle**). The wall thickness of the left coronary artery and ascending aorta measured at 10 points each (n = 10), were 18.4 ± 4.9 µm and 118.5 ± 5.5 µm, respectively, with the lumen area of 8.8 × 10^5^ µm^2^ (**Figure 5C, bottom**), consistent with other 2D measurements.[44, 45] These measurements were obtained from iDISCO-cleared specimens and should be interpreted as values derived from cleared tissue, rather than as absolute anatomical dimensions, due to the potential for clearing-induced tissue deformation.

In addition to myocardial microstructure, this scheme also enables detailed morphological analysis of vasculature. We extracted the coronary arteries by tracing and segmenting the vessels from their aortic origins to distal branches. The binarized left coronary artery (LCA), septal artery (SpA), and right coronary artery (RCA) were digitally traced to identify vessel centerlines, branching points, and local lumen diameters (**Figure 5D** and **Video S5**).[47] Morphological analysis reveals fourth-order vessel branch counts of 411, 141, and 69 for the LCA, SpA, and RCA, respectively, with corresponding mean diameters of 13.4 ± 6.2 µm, 10.4 ± 4.4 µm, and 13.3 ± 4.9 µm. These vessel diameter and branch measurements were obtained from multiple vessel segments within the same representative adult heart and were used to demonstrate the feasibility of AS-DiLS-based vascular segmentation and quantification. Beyond global averaging, spatially resolved analysis of the LCA enables visualization of localized diameter variation ranging from ∼3 µm to 50 µm, capturing fine-detail vessel heterogeneity along its vascular tree (**Figure 5E** and **Video S6**).

Our scheme also enables mesoscale study of the extracellular matrix, a key contributor to cardiac integrity, mechanical behavior, and overall function.[47, 48] Following decellularization and tissue clearing, AS-DiLS was employed for autofluorescence imaging of intact adult rat hearts using three stitched tiles generated from the original 16-bit raw image data (**Figure 5F** and **Video S7**), as well as for imaging of a representative section from the porcine left ventricular wall (**Figure 5G** and **Video S8**). To assess regional variability in the adult rat heart, extracellular volume (ECV) fractions were quantified within three 330 × 330 × 500 µm^3^ volumes in the left ventricular wall (LVW), right ventricular wall (RVW), and interventricular septum (IVS), yielding values of 0.32, 0.33, and 0.26, respectively. This volumetric analysis provides a foundation for characterizing extracellular matrix organization under physiological conditions and offers a framework for future investigations into pathological remodeling processes, such as fibrosis and heart failure.[49, 50]

### 2.6. Topological analysis of vascular network

Specific fluorescence labeling can be effectively integrated into existing imaging and clearing strategies. We investigated the vasculature of intact neonatal mouse hearts using in vivo CD31 labeling. To specifically trace the vascular network in deep tissue, three neonatal mouse hearts at postnatal day 7 (P7) were labeled with CD31 antibody via retro-orbital injection, then fixed in paraformaldehyde at 1-, 2-, and 18-hour post-injection, followed by tissue clearing and imaging for approximately 45 min (**Figure S12** and **Table S1**). AS-DiLS resolves the extensive vascular network, including coronary arteries and microcirculatory networks, in fine detail and contrast throughout the entire heart (**Figures 6A, B** and **Figure S13**), supporting the quantification of endothelial layer thickness in the LCA. We segmented the delineated vascular network in the LCA, RCA, and SpA from their aortic origins to distal branches throughout the intact heart, revealing branching patterns and continuous microvasculature down to seventh-order vessel branches (**Figure 6C** and **Video S9**). For improved visualization, the vessels were segmented as masks that were overlaid on top of the raw data in **Figure 6C** and **Video S9**. Importantly, this method not only reveals vascular density and connectivity, but also facilitates assessment of patency among dense microcirculatory networks, supporting the analysis of vascular injury and remodeling in further studies.[52] To assess microcirculatory features in 3D, we applied Frangi vesselness filtering and filament tracing to reconstruct the capillary network. From each heart, a total of nine representative sections (200 × 200 × 400 µm^3^) were selected from LVW, RVW, and IVS, three from each region, for vessel length density (VLD) quantification (**Figure 6D**). Our statistical analysis of VLD across LVW, RVW, and IVS sections suggests no statistically significant difference among these regions.

**Figure 6.**
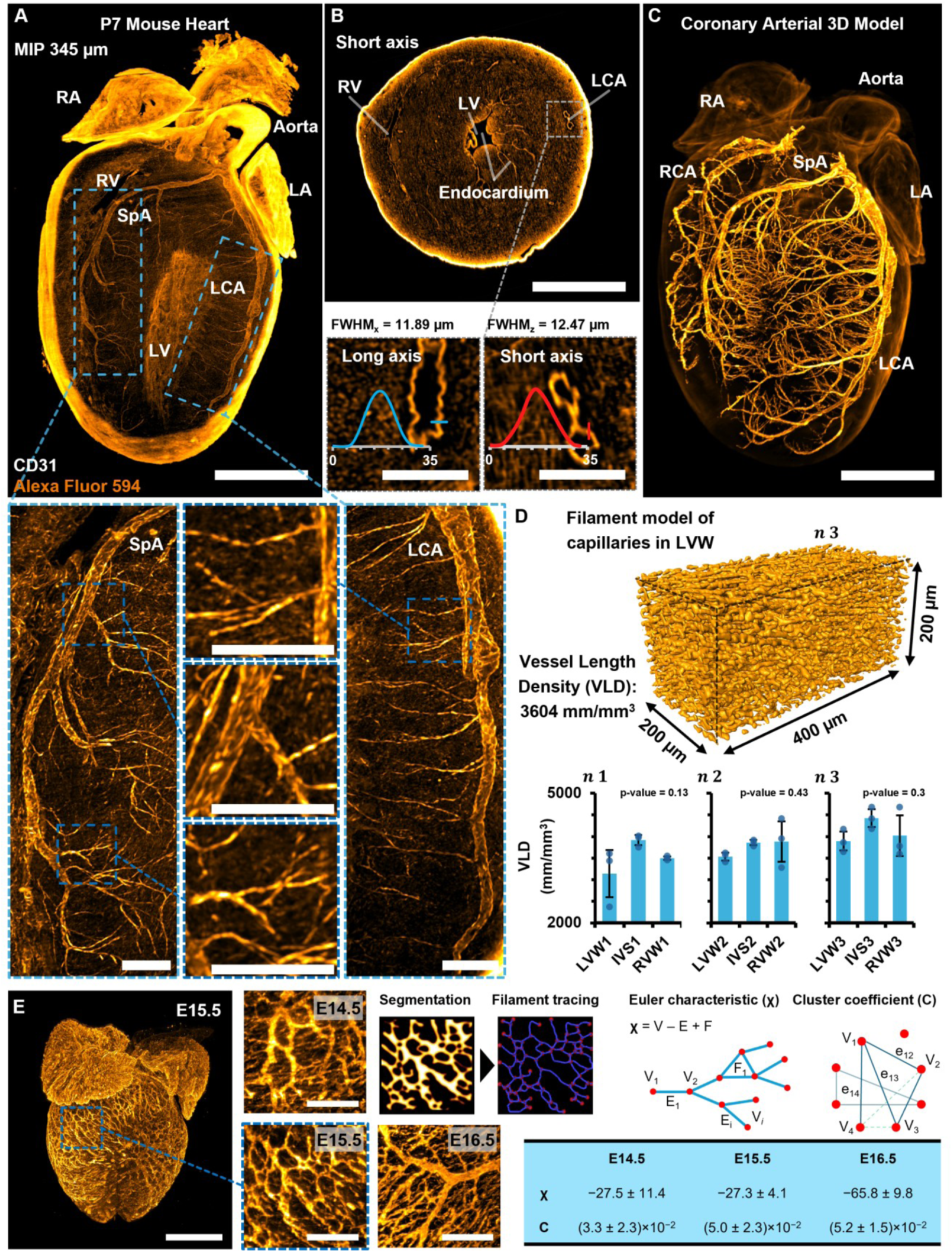
AS-DiLS volumetric immunostaining imaging and topological analysis of the intact neonatal and embryonic mouse heart. **(A)** P7 mouse heart in vivo retro-orbital injection of Alexa Fluor 594 conjugated anti-mouse CD31 to label endothelial cells. MIP image of integrated across 345 µm depth shows uniform vascular labeling and clear delineation of the LCA and SpA from the aorta to distal branches. **(B)** Representative two-dimensional cross-sectional images acquired in the short-axis orientation, illustrating detailed vessel distribution and demonstrating isotropic spatial resolution in both long- and short-axis planes. **(C)** 3D reconstruction of the coronary arterial network, showing the hierarchical branching from main coronary vessels to terminal branches. The LCA and SpA are clearly traced from the aorta throughout the vascular tree. **(D)** Quantitative 3D analysis of capillaries using vessel tracing in Amira. Reconstructions from three CD31-labeled P7 hearts (n = 3) were used to calculate vessel length density (VLD) in three 200 × 200 × 400 µm^3^ sub-volumes, with group statistics reported. **(E)** Volumetric imaging of embryonic hearts at E14.5, E15.5, and E16.5 immunostained with PECAM/Endomucin (Rat-568) and cleared using iDISCO. MIP of E15.5 shows coronary vessel formation. Segmented graphs enabled calculation of Euler characteristics and clustering coefficients across four regions per heart. The centerline and error bars in **(D)** denote the mean and standard deviation, with the associated data points plotted. *P* values were calculated using Student’s *t*-test. RVW, right ventricular wall; LVW, left ventricular wall; IVS, interventricular septum. Scale bars: 1 mm and 200 µm (insets).

In addition to in vivo CD31 labeling, whole-mount immunostaining of vasculature was performed in mouse embryos at E14.5, E15.5, and E16.5, followed by AS-DiLS imaging (**Figure 6E**). This method enables labeling and spatial mapping of the vascular network across the entire heart, allowing imaging through the full 2.2-mm depth of the embryonic heart. The segmented vasculature was converted into graph representations for topological analysis using Euler characteristic (*χ*) and clustering coefficient.[52, 53] Enumeration of graph features, including branching points, vessels, and closed loops of the vascular network, enabled interpretation of network architecture, in which higher *χ* values reflect a sparse, tree-like structure, whereas lower values suggest greater interconnectivity and vascular redundancy. From four representative 260 × 260 × 260 µm^3^ volumes per heart, the mean Euler characteristics were *χ* = −27.5 ± 11.4 for E14.5, *χ* = −27.3 ± 4.1 for E15.5, and *χ* = −65.8 ± 9.8 for E16.5. These regional volumes were analyzed as technical measurements within representative embryonic hearts to demonstrate the feasibility of graph-based topological quantification.

In addition, local vascular connectivity was further assessed using the clustering coefficient (*C_i_*), which quantifies the degree of interconnectivity among neighboring nodes within the vascular graph. For each node, *C_i_* is defined as the ratio of the number of observed connections between its neighboring nodes to the total number of possible connections among those neighbors.[52, 53] This metric provides insight into localized network organization, in which higher *Cᵢ* values suggest denser, more interconnected vascular substructures, while lower values indicate sparser configurations. Analysis of these coefficients across developmental time points revealed changes in microvascular architecture, complementing the global topological trends observed in the Euler characteristic. Global clustering coefficients, calculated from local *C_i_* values across four volumes from three hearts, were *C* = (3.3 ± 2.3) × 10^−2^ for E14.5, *C* = (5.0 ± 2.3) × 10^−2^ for E15.5, and *C* = (5.2 ± 1.5) × 10^−2^ for E16.5, suggesting intense angiogenesis to meet the elevated metabolic demands during this stage of heart development. Collectively, the morphological and topological analyses enabled by AS-DiLS advance our understanding of myocardial microstructure and vasculature in the intact heart with improved speed and space-bandwidth product. This integrated approach holds great promise for facilitating comprehensive and multi-scale assessment of cardiovascular development, pathological remodeling, and therapeutic response, providing an entry point for further investigations into cardiac function and disease progression.

## 3. Discussion

An in-depth understanding of cardiac architecture is crucial for new insights into heart development, remodeling, protection, and aging. Mesoscale volumetric mapping bridges molecular perturbations with organ-level dysfunctions, offering the construction of detailed 3D maps of structural abnormalities across diverse developmental stages and disease progression. However, investigation of centimeter-level patterns and micrometer-level details of cardiac microstructures, including trabeculation, myofibers, valve leaflets, vasculature, and extracellular matrix, remains technically challenging due to photon attenuation within the densely structured myocardial architecture. To address this issue, we have developed the DiLS imaging scheme by creating a time-averaged light-sheet which offers notable flexibility by functioning either as a standalone system or in combination with axial sweeping techniques. The AS-DiLS configuration enables mesoscale data acquisition and analysis of intact ventricles and atria. We have comprehensively analyzed the interplay between laser scanning and rolling shutter sequences and established illumination–detection synchronization that orchestrates their dynamics. By harnessing the intrinsic autofluorescence of myocardial tissue, our method provides high contrast to delineate cardiac architecture and coronary arteries for both global and local assessments. Furthermore, the integration of specific immunolabeling with tissue clearing approaches facilitates full mapping of the vascular network, extending down to the microcirculatory level across the entire heart from embryos to neonates. Complementing this, computational analysis of the imaged cardiac tissue enables morphological and topological assessment of anatomical features, revealing volumetric relationships that are inaccessible through conventional imaging approaches. In combination with computational fluid dynamics or fluid structure interaction, this methodology will further allow investigation of flow capacity and hemodynamics.[54–57]

Based on a dual-actuator configuration integrating beam dithering and axial sweeping, the integration of DiLS with the axially swept approach enables an extended propagation range of up to 12.5 mm, while enhancing imaging speed by over 40% and maintaining uniform optical sectioning and intensity. The principal advantage of AS-DiLS over conventional LSM, DiLS, and ASLM is the coordinated integration of beam dithering with axial sweeping rather than either mechanism alone. Conventional LSM uses a static Gaussian light-sheet and is therefore constrained by the trade-off between light-sheet thickness and confocal region. DiLS addresses this limitation by rapidly dithering the light-sheet waist along the propagation direction, generating a time-averaged illumination profile with an extended effective confocal region. ASLM, in contrast, extends imaging coverage by axially sweeping a static light-sheet in synchrony with the rolling shutter of the sCMOS camera. AS-DiLS combines these strategies by first using dithering to expand the effective confocal region and then sweeping this extended region across the imaging plane in synchrony with rolling-shutter detection. This expanded confocal region increases the number of useful active pixels during each camera exposure, reduces the line-interval constraint during rolling-shutter acquisition, improves tolerance to synchronization mismatch between the swept beam waist and the rolling shutter, and enables faster volumetric imaging while largely preserving optical sectioning across large cardiac specimens. The scheme is also compatible with established stitching and multiview approaches,[58, 59] yet significantly reduces the number of required tiles and viewing perspectives to expand FOV, thereby lowering acquisition and computational demands and supporting near-isotropic resolution of approximately 2.5 µm. Resolution was quantified using 0.53 µm fluorescent beads, which were selected to balance point-emitter approximation with sufficient fluorescence contrast for reliable PSF fitting. The bead diameter was substantially smaller than the measured PSF and approximately four-fold smaller than the expected system resolution under the reported sampling condition, enabling accurate estimation of apparent system resolution.[60] Smaller 0.25 µm beads produced similar resolution measurements (**Figure S7**), but the 0.53 µm beads provided higher fluorescence contrast, allowing shorter exposure times while maintaining sufficient signal-to-background ratio for Gaussian fitting. Therefore, 0.53 µm beads were used for the main PSF characterization experiments. Nevertheless, the experimentally measured FWHM represents the convolution of the optical PSF with the finite bead diameter and may also be influenced by bead-to-bead variability, residual aggregation, local embedding heterogeneity, and fitting uncertainty. Accordingly, these measurements should be interpreted as apparent experimental resolution values rather than ideal diffraction-limited PSF estimates. Although CV provides a useful descriptive measure of relative axial-resolution variability, its interpretation depends on the spatial domain over which measurements are collected. In this study, LSM and DiLS were evaluated within smaller usable confocal regions, whereas AS-DiLS was evaluated across a substantially larger swept imaging field. Therefore, the CV analysis supports the assessment of resolution uniformity within each configuration but should not be interpreted as a direct statistical comparison across identical sampling domains. Because each configuration produced a single CV value, bootstrap resampling was used to estimate the uncertainty associated with each CV estimate and to generate a 95% confidence interval, thereby providing an uncertainty range that supports the reliability of the reported CV values within their corresponding analyzed regions. Notably, uniform optical sectioning and intensity in the time-averaged light-sheet are achieved by synchronizing rapid laser dithering with line interval delays within a single camera exposure. The dithering frequency can be configured as an integer multiple of the camera’s readout rate to ensure temporal alignment, and our customized script in MATLAB is available at Zenodo.[39] As an external validation of generalizability, we also implemented AS-DiLS on an independently configured benchtop-mesoSPIM platform[61] in the Dean laboratory, operated with separate control software.[62] Under matched acquisition conditions, AS-DiLS produced higher signal intensity than ASLM, consistent with the performance trends reported here (**Figure S21**). Notably, in contrast to closed-loop actuators such as piezo stages[27] or VC scanners[25], the ETL operates in an open-loop configuration, which can introduce axial positioning errors at higher imaging frequencies (e.g., > 2 Hz in our system). Such errors compromise the tight synchronization required in ASLM, where precise overlap between the swept confocal region and the rolling shutter is essential. In contrast, the extended confocal region in DiLS provides greater tolerance to such mismatches by increasing the effective overlap between the swept beam and the rolling shutter, thereby mitigating synchronization errors and enabling a thinner effective light-sheet at higher imaging frequencies. In our implementation, the frequency was set to twice the reciprocal of the exposure time, constrained by the operational limits of the VC scanner. Although dithering extends the confocal region, it is accompanied by a modest broadening of the effective light-sheet. Therefore, AS-DiLS should be interpreted as improving the balance among confocal-region length, imaging speed, and axial resolution, rather than eliminating this optical trade-off entirely. Future integration of high-speed scanners with closed-loop feedback control holds promise for further improving imaging speed and throughput. Moreover, optimizing the dithering frequency enables a tunable balance between light-sheet thickness and scanning velocity. The resulting expansion of the confocal region and improved data acquisition rate, modulated by dithering frequency, make the DiLS scheme highly adaptable for a broad range of imaging applications. Beyond dithering frequency optimization, the AS-DiLS scheme could be further improved using multiple foci generated by other devices like digital micro-mirrors,[63] spatial light modulator,[64] and multilayer beam splitter.[65] AS-DiLS can also be combined with dual-sided illumination to reduce shadowing and extend effective illumination length, although this configuration was excluded from the current study owing to the technical challenges of maintaining co-alignment of two light-sheets across large fields of view, which can introduce imaging artifacts.[29] Additionally, to correct for the Gaussian distribution of laser intensity along the light-sheet height, a galvanometer-based laser scanning or Gaussian blur-based post-processing could be employed to achieve evenly distributed illumination, thereby improving uniformity across the imaging volume.

The current limitation in pixel size on the sCMOS camera presents a fundamental challenge for digital sampling in mesoscale imaging. Although the theoretical optical resolution of our system approaches ∼1 µm, it is constrained by a maximal digital sampling rate of 1 µm under 6.3× magnification. In contrast to customized objective lens designs, our subvoxel sampling techniques[41, 64] offer a universal solution for resolution enhancement through sequential subvoxel shifting mechanisms, thereby increasing the system’s space-bandwidth product through widely used optical components. In terms of stripe artifacts, commonly caused by photon attenuation in LSM, AS-DiLS allows for minimal effects through dynamic dithering and sweeping (**Figure 4E** and **Figure S11**). To mitigate the decline in SBR and CNR observed at the distal end of light propagation, multi-directional illumination strategies could be incorporated. These approaches have the potential to collectively improve image fidelity and extended imaging depth, making AS-DiLS a robust platform for mesoscale volumetric mapping of intricate microstructure across multiple organ systems under physiological and pathophysiological conditions.

Another limitation of the current study is that quantitative anatomical measurements were performed on cleared specimens. Although iDISCO-based clearing enables high-resolution volumetric imaging of intact cardiac tissue, processing steps such as dehydration and delipidation may lead to tissue deformation, such as shrinkage[32] Despite this potential limitation, relative comparisons among experimental groups remain valuable when all samples are processed using the same tissue-clearing protocol and imaging workflow, and analytical framework.[19] However, measurements such as myocardial fiber thickness, vessel-wall thickness, lumen area, and extracellular matrix volume fraction should be interpreted as values obtained from cleared tissue rather than absolute anatomical dimensions. Future studies requiring absolute anatomical measurements should incorporate clearing-related correction factors or validation using complementary imaging or histological approaches.

We anticipate that these implementations, built on the dual-actuator AS-DiLS framework, will expand efficiency and broaden the utility of the AS-DiLS framework, extending its impact from fundamental research to translational discoveries in cardiovascular research and beyond. In parallel, advances in oblique and open-top imaging system designs,[65–69] deep learning tools,[70, 71] and extended reality[72–75] offer unprecedented opportunities for high-dimensional feature extraction and interpretation of cellular and subcellular structures within intact organs and cultured systems. The continued integration of these technological innovations with extensively established animal models and human-derived cell cultures is poised to deepen mechanistic understanding, facilitate targeted therapeutic strategies, and ultimately bridge transformative discovery with human-centered clinical investigations.

## 4. Materials and Methods

### 4.1. System design

This in-house system integrates four configurations: conventional LSM, ASLM, DiLS, and AS-DiLS (**Figure S1**). In conventional LSM mode, a static Gaussian light-sheet was used without dithering or axial sweeping. In DiLS mode, the light-sheet waist was rapidly displaced along the propagation direction using the VC scanner, generating a time-averaged beam profile with an extended effective confocal region. In ASLM mode, a static light-sheet was axially swept using ETL and synchronized with the rolling shutter of the sCMOS camera. In AS-DiLS mode, the VC scanner and ETL were operated together so that the dithered, extended confocal region was swept across the imaging plane in synchrony with rolling-shutter detection. It consists of three continuous-wave diode-pumped solid-state lasers with wavelengths of 473 nm, 532 nm, and 589 nm (LRS-0473-PFM-00100-05, LRS-0532-PFM-00100-05, LRS-0589-GFF-00100-05, Laserglow Technologies). The diameter of laser beams was controlled using a 10× achromatic beam expander (GBE10-A, Thorlabs) and an iris (ID20/M, Thorlabs). A remote focusing design including a VC flexure scanner (VCFL35, Thorlabs), a half-wave plate (AHWP3, Bolder Vision Optik), a polarizing beam splitter (CCM1-PBS251/M, Thorlabs), a quarter-wave plate (AQWP3, Bolder Vision Optik), and an objective lens (Plan Fluor 10×/0.3 NA, Nikon)[36, 37] was implemented to generate the DiLS in the illumination path. The beam reflected from the VC scanner was focused using a plano-convex cylindrical lens (*f* = 50 mm, ACY254-050-A, Thorlabs) to form a light-sheet. The ETL (EL-16-40-TC, Optotune) was vertically mounted on the focal plane of the cylindrical lens to axially sweep the dithered beam, serving as the second scanner in AS-DiLS. A series of achromatic doublets (*f_1_* = 100 mm, *f_2_* = 60 mm, AC254-100-A, AC254-060-A, Thorlabs) and the objective lens (Plan Fluor 10×/0.3 NA, Nikon) were used to relay and reshape the beam into a vertical light-sheet in the imaging plane.

The detection module was built upon our established LSM system[38] that includes an objective lens (1×/0.25 NA, MV PLAPO, Olympus), a zoom body (0.63–6.3×, MVX-ZB10, Olympus), a filter wheel (LB10-W32, Sutter Instrument), a set of filters (ET525-30, ET585-40, ET645-75, Chroma), a tube lens (MVX10, Olympus), and an sCMOS camera (Flash 4.0 v3, Hamamatsu). The entire detection module was mounted on a high-precision translational stage (C-884.4DC, Physik Instrumente) to align the detection focus with the light-sheet.

A custom-designed chamber was 3D-printed to isolate the tissue clearing solution from the detection lens, providing a uniform refractive index environment, leveraging the extended depth of field, and ensuring compatibility with multiple clearing protocols.[40] For precise positioning, we used a 4D motorized stage that includes 3D translational movement (L-406.20DG10, L-406.40DG10, M-403.4PD, controller C-884, Physik Instrumente) and 1D rotation (1474 stepper motor, Pololu).

### 4.2. System control

A customized control algorithm developed in LabVIEW (2017 SP1, National Instruments), along with NI VAS (20.0), compactRIO (20.0), FlexRIO (20.0), R-series (20.0), Serial (20.0), DAQmx (20.1), and a DAQ card (PCIe-6363, National Instruments), was utilized to enable the system synchronization. To enable smooth translation of the VC scanner, a filtered triangular waveform was generated via the NI card, amplified through a TI module (OPA548EVM, Evaluation Module, Texas Instruments), and powered by a 3-channel supply unit (T3PS43203, Teledyne LeCroy) (**Figures S14, 15**). Axial sweeping was achieved at a constant velocity using an ETL driven by an independent triangular trigger ranging from 0.25 to 4 Hz, with a 2.5 V offset and an amplitude of 0 to 2.5 V (**Figure S16**). We diagnosed synchronization errors by rotating the camera 90°, aligning the rolling shutter vertically against the light-sheet sweep. Ideal synchronization produced a diagonal line, indicating precise matching of ETL scanning and shutter activation (**Figures S17, 18**). Image acquisition on the sCMOS camera was triggered by an edge signal from the NI card, and an empirically determined 8 ms phase delay was introduced to synchronize the ETL with the camera (**Figure S19**). In AS-DiLS, the exposure time serves as the primary control parameter, from which the line interval and the corresponding dithering and sweeping frequencies are derived. The line interval also governs the rolling shutter speed, ensuring precise temporal coordination between active pixel rows and the beam waist (**Notes S2-3**).

### 4.3. Benchtop-mesoSPIM

External validation experiments were performed on a slightly modified benchtop-mesoSPIM platform with single-sided illumination. Samples were imaged in a 50 mm cuvette (Hellma, Z805750) using a macro objective lens (4×/0.28 NA, XLFLUOR, Olympus) with a working distance of 29.5 mm. Excitation was provided by a laser combiner (L4Cc-CSB-1211-488-561-638-300, Oxxius), with the 638 nm channel filtered using a 20 nm bandpass cleanup filter. The microscope was equipped with motorized translation stages for X, Y, Z, focus, and rotation (Applied Scientific Instrumentation), a six-position filter wheel (FW-1000, Applied Scientific Instrumentation), a controller (Tiger Controller, Applied Scientific Instrumentation), galvanometric scanners (QS15X-AG, Thorlabs), and a sCMOS camera (ORCA-Flash4.0 V3, Hamamatsu). Axially swept operation was enabled using ETL (EL-16-40-TC-VIS-5D, Optotune) driven by a lens controller (EL-E-4i, Optotune). The system was operated using open-source Navigate software, with synchronized analog and digital control provided by a data acquisition card (PCIe-6723, National Instruments).

### 4.4. Image rendering

Image stitching was performed to reconstruct the adult heart dataset. Individual 16-bit TIFF files were imported into the BigStitcher plugin in Fiji (version 1.54f) for stitching and fusion.[76–78] Datasets exceeding 5 GB were downsampled using 2× binning and converted into 8-bit images in Fiji for 3D rendering in Amira (2021.2; Thermo Fisher Scientific).[47]

### 4.5. Simulation of static and dithered light-sheet

A static Gaussian light-sheet was synthesized using the Fourier transform and angular spectrum propagation methods to obtain its cross-sectional intensity profile. In the spatial frequency domain, the illumination beam was described as:

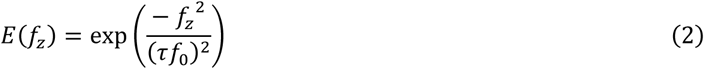

where *τ* is a shape parameter that controls beam thickness (*τ* = 0.08), and *f_0_* = *n*/*λ* is the characteristic spatial frequency determined by the refractive index (*n* = 1.33) and wavelength (*λ* = 532 nm). The inverse Fourier transform of *E(f_z_)* was used to obtain the axial field distribution at the focal plane, corresponding to the light-sheet waist. The field was propagated using the angular spectrum method to obtain its evolution along the propagation axis. The light-sheet intensity distribution was then obtained as the squared magnitude of the complex field. In the dithered configuration, the Gaussian light-sheet was rapidly dithered along the propagation axis, and its effective profile was defined by the time-averaged intensity. At each spatial position, the time-averaged intensity was calculated by integrating the instantaneous field intensity over a dithering period (*Δt*). The total displacement during one cycle was *v* × *Δt*, corresponding to twice the dithering amplitude. For both static and dithered light-sheet, the thickness was defined as the full width at half maximum (FWHM) of the intensity profile at the waist.

### 4.6. Point spread function calibration

Fluorescent beads (0.53 µm, FP-0556-2, Spherotech) were diluted to a concentration of 1:1×10^5^ in a 1% low-melting point agarose solution (16520050, Thermo Fisher Scientific) vortexed at 4500 rpm for 2 min using a vortex mixer (BenchMate Mini Vortex Mixer VM-M, Oxford Lab Products). They were mounted in a fluorinated ethylene propylene tube and placed in a chamber filled with double-deionized water for imaging. PSF measurements were performed using raw 16-bit image datasets. The intensity profile of each bead was fitted with a Gaussian distribution in MATLAB, and FWHM was calculated to estimate lateral and axial resolution.[81]

### 4.7. Beam profile test

A 50 µM fluorescein solution was utilized to record the beam profile. It was prepared by dissolving 3 mg of fluorescein in 180 mL of deionized water, followed by mixing with 20 mL of 10× phosphate buffered saline (PBS) in the imaging chamber. To capture the beam profile, the cylindrical lens was replaced with an achromatic doublet lens of identical focal length (AC254-050-A, Thorlabs).[81]

### 4.8. Animal specimens

All animal protocols, experiments, and housing described in this manuscript were approved by the Institutional Animal Care and Use Committee (IACUC #20-07 and #21-03) of the University of Texas at Dallas and other institutions from which the samples were obtained.

### 4.9. Tissue clearing

All hearts were cleared under the iDISCO protocol.[31, 42] Fixation was performed in 4% paraformaldehyde, adult hearts for 18 hours, embryonic hearts for 2 hours, and three neonatal hearts for 1, 2, and 18 hours, respectively. Following fixation, hearts were rinsed three times in 1× PBS for 30 min each. These samples were then dehydrated at room temperature by agitating in methanol with increasing concentrations (20%, 40%, 60%, 80%, and 100%), each for 1 hour. Delipidation was carried out by overnight immersion in dichloromethane at room temperature, followed by two 30-min washes in methanol. Hearts were bleached in 5% H_2_O_2_ in methanol overnight at 4°C. These samples were rehydrated through a reverse methanol gradient (100%, 80%, 60%, 40%, and 20%) and finally in 1× PBS at room temperature, each for 1 hour. To enhance permeability, hearts were washed twice in 0.2% Triton X-100 in 1× PBS at room temperature for 1 hour. Embryonic and neonatal hearts were ready for optional immunolabeling. Subsequently, hearts were embedded in a 1% agarose prepared in 1× PBS. They were dehydrated again in methanol with increasing concentrations (20% to 100%) for 1 hour each, followed by immersion in a 66% dichloromethane / 33% methanol mixture at room temperature for 3 hours to complete lipid removal and allow the samples to sink. Finally, hearts were submerged in dibenzyl ether until fully transparent.

### 4.10. In vivo CD31 immunolabeling

Neonatal mice P7 were anesthetized by isoflurane. Each mouse then received a retro-orbital intravenous injection[54, 80] of 20 µL rat anti-mouse CD31 antibody conjugated to Alexa Fluor 594 (102520, BioLegend), diluted 1:5 in 1× PBS. After the injection, the mice were returned to their mother for 30 min prior to euthanasia. The hearts were then perfused with 2 mL of 4°C 1× PBS through the left ventricle. Hearts were harvested and placed into 4°C 1× PBS and gently compressed with flat-tip forceps to remove residual blood. Following the iDISCO protocol, the hearts were blocked in a solution containing 10% dimethyl sulfoxide (DMSO) and 6% normal donkey serum (NDS; 017-000-121, Jackson ImmunoResearch) in 0.2% Triton X-100 for 2 days at 37°C for immunolabeling. Alexa Fluor 594-conjugated secondary antibodies (712-585-153, Jackson ImmunoResearch) were diluted 1:1 in 1× PBS containing 3% NDS, 0.2% Tween-20, and 0.1% heparin and the hearts were immersed for 7 days at 37°C. This secondary antibody incubation was used to amplify the signal from the retained rat anti-mouse CD31 antibody after fixation, dehydration, delipidation, and bleaching, rather than to introduce an additional vascular marker. The hearts were then washed every 1 hour in 1× PBS containing 0.2% Tween-20 and 0.1% heparin (H4784-1G, Sigma) five times, followed by an overnight wash. Finally, the samples were cleared using the remaining steps of the iDISCO protocol.

### 4.11. Immunolabeling in the embryonic heart

Embryonic mouse hearts were immunolabeled following iDISCO protocol. 2.5 µL of 1% biotin-conjugated Cholera Toxin Subunit B was injected slowly. Hearts were incubated in a permeabilization solution containing 0.2% Tween-20 and 0.1% heparin in 1× PBS for 2 days at 37°C in a water bath. Following permeabilization, samples were blocked in a solution of 20% DMSO, 0.2% glycine, and 0.2% Triton X-100 for 2 days at 37°C. A second blocking step was performed in 10% DMSO and 6% serum in 0.2% Triton X-100 for an additional 2 days at 37°C. For primary antibody incubation, PECAM and Endomucin were diluted in 1× PBS containing 5% DMSO, 3% NDS, 0.2% Tween-20, and 0.1% heparin, and incubated with the samples for 7 days at 37°C. Following primary labeling, hearts were washed five times in PBS containing 0.2% Tween-20 and 0.1% heparin for 1 hour each, followed by an overnight wash. Secondary antibody incubation was carried out using Alexa Fluor 568, diluted in PBS with 3% NDS, 0.2% Tween-20, and 0.1% heparin. Hearts were incubated for 7 days at 37°C. Post-secondary incubation, hearts were washed five times in PBS containing 0.2% Tween-20 and 0.1% heparin for 1 hour each, followed by an overnight wash. Finally, immunolabeled hearts were cleared using the remaining steps of the iDISCO protocol.

### 4.12. Vessel segmentation, tracing, and quantification

Manual segmentation of the coronary artery lumen in the adult mouse heart was performed on raw 16-bit image datasets based on the geometry and intensity difference between the vessel wall and the lumen area. Segmentation started at the origin of the left, right, and septal coronary arteries at the aorta and continued to the fully resolved vessel branches, using 2D cross-sectional images. The aorta was used as the main anatomical branch point to guide manual segmentation of the coronary artery branches. The coronary autofluorescence signal was continuous in the analyzed dataset, allowing vessel branches to be followed across adjacent optical sections. To quantify local vessel morphology, filament tracing was conducted to convert the segmented volume into a graph-based representation. The centerlines were extracted to compute geometric features, including the number of vessel branches (edges), branching points (nodes), and local radii. The radius along the centerline was calculated based on the minimum distance to the vessel boundary, assuming a circular cross-section. The spatial variations of local lumen radius throughout the coronary network were visualized in a 3D color-coded map, where the smallest radii were shown in blue and the largest radii in red. VLD was calculated by cropping 200 × 200 × 400 µm sub-volumes from whole-heart image stacks. The cropped 16-bit image stacks were denoised by applying Gaussian smoothing (*σ* = 1 µm), followed by background subtraction with a rolling ball radius of 20 µm. A Frangi vesselness filter[81, 82] was applied to enhance vessel contrast and improve continuity, particularly for capillaries. The processed image stack was used for subsequent analyses. For embryonic mouse hearts, 16-bit image stacks were processed using Gaussian smoothing (*σ* = 1 µm) and background subtraction with a rolling ball radius of 20 µm, followed by filament tracing in Amira to generate vascular graphs for Euler characteristic and clustering coefficient calculations.

### 4.13. Extracellular matrix sample preparation

Decellularized extracellular matrix of rat and porcine hearts was prepared following adapted protocols.[83, 84] The left ventricle of adult porcine hearts (Animal Technologies, Inc.) was dissected, and excess vasculature, adipose tissue, and connective tissue were removed. The isolated myocardium was sectioned into ∼1 mm thick sections and rinsed extensively with deionized water to remove residual blood. Tissues were decellularized in 1% sodium dodecyl sulfate (SDS) solution for 4 days at room temperature, with daily replacement of fresh SDS, followed by an overnight rinse in deionized water to remove residual detergent. The rat heart was harvested from a euthanized Sprague-Dawley rat via thoracotomy. After removing adipose and connective tissue, the heart was decellularized in 1% SDS solution for 3 weeks at room temperature. Fresh SDS was replaced daily during the first week and every other day thereafter to ensure effective detergent exchange. To enhance tissue permeability, 1% SDS was injected twice daily through the aorta using a syringe. Upon completion of detergent treatment, hearts were rinsed thoroughly in excess deionized water for 24 hours to remove residual SDS. The decellularized tissues were dehydrated and rehydrated using the iDISCO protocol to prepare them for imaging.

### 4.14. Coefficient of variation

The CV, defined as the ratio of the standard deviation (*σ*) to the mean (*μ*) of the measured FWHM, was calculated to assess the uniformity of resolution:

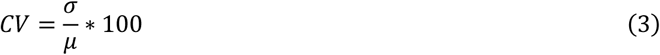

CV values less than 10% were considered acceptable uniformity in this study. As each imaging configuration produced a single CV value, bootstrap resampling was used to estimate the uncertainty associated with each reported CV. For each configuration, 10,000 bootstrap samples were generated by resampling the measured axial FWHM values with replacement using the same sample size as the original dataset. The CV was recalculated for each bootstrap sample, and the resulting bootstrap CV distribution was used to report the 95% confidence interval. This approach provides an uncertainty range around each CV value and supports the reliability of the reported CV estimates.

### 4.15. Contrast-to-noise ratio and signal-to-background ratio

The CNR and SBR were calculated from raw 16-bit image datasets as follows, respectively:[87]

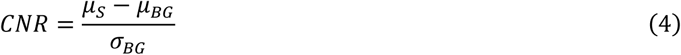

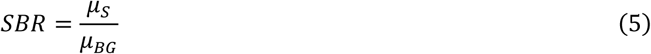

where *μ_S_* is the mean intensity of fluorescence measured within the region of interest, *μ_BG_* is the mean intensity of the background, and *σ_BG_* indicates the standard deviation of the background intensity. Higher CNR values indicate a clearer distinction between the fluorescence signal and background, whereas higher SBR values reflect a greater proportion of fluorescence photons relative to background photons.

### 4.16. Fourier transform analysis

Fourier transform analysis was performed using raw 16-bit images. Image quality improvement from LSM to AS-DiLS was quantified by estimating the ratio of OTF support radii between two images from the same region.[44] Fourier power spectra were obtained by applying the Fast Fourier Transform to both the AS-DiLS image and LSM image. Edge effects were removed by applying a Hanning window, and signal power was isolated using background power subtraction, both ensuring accurate and consistent OTF support radii estimation. The cut-off radius used to quantify image quality improvement was determined through radial projections of the Fourier spectra, determined by the maximum projection length before falling below a power threshold of 1%:

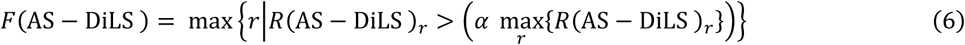

Repeating these steps for LSM image, then comparing the ratio of the OTF support radii (*F*(AS-DiLS) / *F*(LSM)) quantifies relative image improvement, where values greater than 1 indicate enhanced image quality and resolution in AS-DiLS image. This method supplements spatial analysis by providing insights into the frequency relationships affecting imaging system parameters, particularly in complex biological tissues, where point-emitter-based OTF measurements cannot be implemented. To eliminate the effect of horizontal stripes in LSM imaging during Fourier transform analysis, artifacts observed along the k_y_-axis in the Fourier domain were removed using frequency-domain filtering.

### 4.17. Topological analysis of the vascular network

The Euler characteristic (*χ*), a global topological metric reflecting the balance between branching, connectivity, and looping of the network,[53] was used to quantify the overall shape and connectivity of the vasculature, and is defined as:

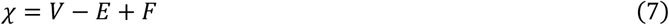

where *V* is the number of branching points or endpoints, *E* is the number of vessel segments, and *F* is the number of closed-loop vessels. A high value indicates a simpler, tree-like structure with less interconnectivity, whereas a low value suggests a more complex and interconnected vascular architecture.[88]

To assess local connectivity within the vascular network, the clustering coefficient was calculated as a measure of the tendency for branching points to form interconnected neighborhoods. For each node, the local clustering coefficient *C_i_* was computed based on the number of actual connections between its neighboring nodes relative to the total possible connections among them, and is defined as:

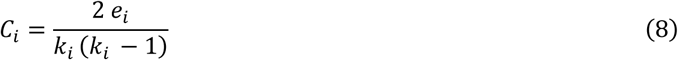

where *e_i_* is the number of edges between the *k_i_* neighbors of node *i*, and *k_i_*(*k_i_* -1)/2 represents the total number of possible connections among neighbors of node *i.*[89] A higher clustering coefficient indicates a more interconnected local neighborhood.

The global clustering coefficient, defined as the average of local coefficients across all branching points, was calculated as:

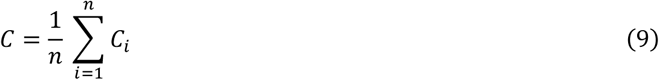

where *C* is the global clustering coefficient, and *n* is the number of branching points.

### 4.18. Structural similarity index measure

The SSIM was employed to assess the consistency of structural information during the axial sweeping across the imaging plane using raw 16-bit image datasets, and is defined as:[90]

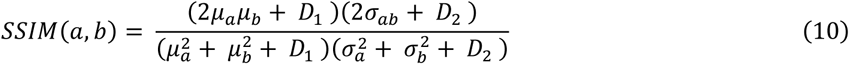

where *μ_a_*and *μ_b_* are the mean intensities of images *a* and *b*; *σ*_a_ ^2^ and *σ* ^2^ are the variances of images *a* and *b*; *σ_ab_* is the covariance between two images; *D_1_*and *D_2_* are the variables used to stabilize the division with a small denominator, and are determined as: *D_1_* = (0.01 × *L*)^2^ , *D_2_* = (0.03 × *L*)^2^, where *L* = 2^16^-1 for 16-bit images.

### 4.19. Statistical analysis

For biological experiments, individual specimens were considered the biological units. Measurements obtained from multiple vessel branches, image regions, sub-volumes, or voxels within the same specimen were treated as technical or nested measurements and were not interpreted as independent biological replicates. Bead-based PSF measurements were used for optical system characterization and were therefore treated as technical resolution measurements, consistent with standard practice in light-sheet microscopy. Vessel-branch, VLD, and image-quality measurements were used to demonstrate the feasibility of AS-DiLS for quantitative analysis of intact cardiac specimens, rather than to draw powered biological conclusions across experimental groups. Accordingly, these measurements were summarized within their corresponding specimen, region, or imaging condition and interpreted in the context of technical performance and analytical feasibility. All data are presented as mean ± standard deviation unless otherwise specified. Confidence intervals are reported where applicable. For CV analysis, 95% confidence intervals were calculated using bootstrap resampling. For datasets that deviated from normality or did not satisfy parametric assumptions, data were additionally summarized as median and interquartile range (IQR) where appropriate. Box plots display the median and IQR, with whiskers extending to 1.5 × IQR, and individual points represent independent measurements. All statistical tests were two-sided. Comparisons among multiple groups were performed using one-way analysis of variance (ANOVA). For pairwise comparisons between independent groups, unpaired two-tailed Student’s t-tests were applied when parametric assumptions were satisfied; otherwise, the Wilcoxon rank-sum test was used. Differences were considered statistically significant at *p* < 0.05.

### 4.20. Workstations

Images were collected by a dedicated workstation equipped with a processor (Xeon W-2245, 3.9 GHz, 8 Core, Intel), a graphics card (Quadro RTX 4000, NVIDIA), and 256 GB of RAM. The motherboard houses several PCIe cards, including 2 CameraLink frame grabbers for streaming images from the camera, and a DAQ card (PCIe-6363, National Instruments) for generating analog output voltages. For image post-processing, computational analysis, and rendering, a workstation equipped with a processor (Xeon Gold 6258R, 2.69 GHz, 56 Core, Intel), two graphics cards (RTX A6000 GPU, NVIDIA), and 770 GB of RAM was used.

## Acknowledgments

We express our gratitude to Dr. Fenghua Shi for valuable discussions on simulation at UT Dallas. We also appreciate the constructive comments and technical support provided by the D-Incubator members at UT Dallas. This work was financially supported by the National Institutes of Health grants R00HL148493 (Y.D.), R01HL162635 (Y.D.), and RM1GM145399 (K.M.D.), the National Science Foundation grant 2503230 (Y.D.), and the Cecil H. and Ida Green Professorship in Systems Biology Science (Y.D.).

## Conflict of Interest

K.M.D. has a patent covering ASLM (US10989661), consultancy agreements with 3i, Inc (Denver, CO, USA), and an ownership interest in Discovering Imaging Systems, LLC. The remaining authors declare no conflict of interest.

## Author contributions

**Conceptualization:** MA, YD **Methodology:** MA, AS, ESB, XZ, YD **Software:** MA, AS, ESB, RH, YD

**Validation:** MA, AS, ESB, XZ, RH, YL, SAW, SZ, JL, MW, DT, YD

**Formal analysis:** MA, AS, ESB, XZ, RH, HH, JY, YD

**Investigation:** MA, YL, SAW, JL, YH, MW, DT, DS, YD

**Resources:** MA, YL, SAW, SZ, YW, HL, KMD, JL, YH, MW, DT, DS, YD

**Data curation:** MA, AS, ESB, XZ, RH, HH, YD

**Writing – original draft:** MA, AS, ESB, YL, YW, YH, MW, YD **Writing – review & editing:** MA, AS, ESB, XZ, JC, SAW, HH, DT, YD **Visualization:** MA, AS, ESB, JY, YD

**Supervision:** YD

**Project administration:** YD

**Funding acquisition:** YD

## Data and materials availability

MATLAB codes, and software for calculating AS-DiLS are publicly available at Zenodo.[39] Additional information required to reanalyze the data reported in this paper is available from the corresponding author upon request.

## Funding

National Institutes of Health grant R00HL148493 (Y.D.) National Institutes of Health grant R01HL162635 (Y.D.) National Science Foundation grant 2503230 (Y.D.)

Cecil H. and Ida Green Professorship in Systems Biology Science (Y.D.) National Institutes of Health grant RM1GM145399 (K.M.D.)

## Supplementary Materials

**The file includes:**

Supplementary Notes 1 to 3

Supplementary Figures 1 to 21

Supplementary Table 1

Legends for Supplementary Video 1 to 9 References

## Other Supplementary Material for this manuscript includes the following

Supplementary Videos 1 to 9

## Notes

### Competing Interest Statement

The authors have declared no competing interest.

### Summary of Updates

We have updated the title, abstract, and authorship of this manuscript. The conclusion remains the same as previous versions.

